# Tracing SARS-CoV-2 Evolution in Algeria: Insights from 2020–2023

**DOI:** 10.1101/2025.05.28.656620

**Authors:** Fatima Ezzohra Ezahedi, Fawzi Derrar, Ágota Ábrahám, Safia Zeghbib

**Author notes:** **Institutions Where the Work Was Performed:** National Laboratory of Virology, Szentágothai Research Centre, University of Pécs, Pécs, Hungary Pasteur Institute of Algeria. **Corresponding Author:** Safia zeghbib University of Pécs, National laboratory of virology Faculty of Sciences Pécs, Ifjúság útja, 7624.

## Abstract

SARS-CoV-2 continues to circulate globally, leading to the emergence of new viral sublineages. This study examined 449 full-length genomic sequences from Algeria (March 2020 to May 2023) to explore SARS-CoV-2 evolution in this region. Our analysis revealed multiple introductions of viral strains, which were marked by significant genomic changes over time. Algeria-specific mutations were identified, providing insights into regional adaptations and evolutionary dynamics. Furthermore, a statistically significant recombination event was detected using the RDP 5 software. EPI_ISL_15920753 was recombinant, whereas EPI_ISL_15790700 and EPI_ISL_12156732 constituted the major and minor parents, respectively. Notably, haplotype analysis revealed 179 distinct haplotypes, including a highly connected node (H91) associated with a superspreading event later in the pandemic (July–September 2022), suggesting rapid transmission likely driven by mobile individuals and large gatherings. These findings emphasize the ongoing evolution of SARS-CoV-2 in local settings and highlight the importance of genomic surveillance for tracking emerging mutations and their potential impact.

**Importance:** Bridging a critical gap in North African genomic surveillance, this study presents the first comprehensive evolutionary analysis of SARS-CoV-2 in Algeria using a significantly larger dataset of 449 sequences, a substantial increase from the only previous study of 29 genomes conducted at the onset of the pandemic. Our results provide essential insights into how the virus was introduced, disseminated, and adapted within the country, thereby guiding regional public health initiatives and supplying important data for global comprehension of the evolution of SARS-CoV-2.

## Introduction

Despite public health measures to effectively manage the SARS-CoV-2 pandemic and the success of vaccination in easing restrictions, the persistent threat posed by emerging variants remains a significant challenge. The substantial evolutionary adaptability of this virus is evident from the proliferation of variants characterized by increased transmissibility. Concerns persist regarding the potential emergence of novel variants capable of evading immunity acquired through vaccination or previous infection^1^.

Coronaviruses are a diverse group of viruses that can cause symptoms ranging from mild, such as the common cold, to severe illnesses, such as severe acute respiratory syndrome (SARS). Studies have shown that asymptomatic individuals can carry viral loads similar to those of symptomatic individuals and transmit the virus, complicating efforts to control it ^2^. The SARS-CoV-2 genome, which is approximately 30,000 base pairs long, encodes four structural proteins and 16 nonstructural proteins. The genes encoding these proteins included *ORF1a, ORF1b, S, ORF3a, E, M, ORF6, ORF7a, ORF7b, ORF8, N,* and *ORF10*. *ORF1ab* encodes NSP1, which binds to the 40S ribosomal subunit and blocks host mRNA entry ^3^. NSP2 and NSP3 act as transmembrane helices, with NSP2 mutations enhancing contagiousness and NSP3 mutations potentially differentiating COVID-19 from SARS ^4^. Moreover, NSP4 facilitates extracellular mtDNA release by inducing the formation of outer membrane macropores and inner membrane vesicles, together with Orf9 ^5^. NSP5 induces replication by activating mitochondrial antiviral signaling. Additionally, it potently inhibits interferon (IFN) induction by double-stranded RNA (dsRNA) in an enzyme-dependent manner ^6^. NSP6 oligomerizes and forms an amphipathic helix that joins the ER membrane and establishes connectors ^7^. NSP7 and NSP8 are accessory factors of RdRP; they serve as cofactors for NSP12, stabilizing it, while NSP8 acts as a primase-like enzyme ^8^. NSP10 joins NSP16 in a complex to methylate the first nucleotide of a capped RNA strand at the 2’-O position ^9^. Likewise, NSP10 forms a complex with NSP14 to provide a proofreading function owing to the exoribonuclease activity of NSP14 ^10^. Additionally, NSP13 was found to inhibit IFN-I pathway activity by forming a complex with TBK1 and p62 ^11^. Furthermore, NSP15 was found to be a uridine-specific endoribonuclease that helps the virus evade triggering an innate immune response ^12^.

The first confirmed case in Algeria was reported on February 25, 2020. Previous studies analyzing 95 genomes, including 29 from Algeria, revealed multiple virus introductions despite lockdown. Moreover, an individual deleterious mutation, A23T, in ORF3a was identified in one Algerian sequence ^13^. However, the limited sample size has constrained findings on viral evolution and spread within the country. To address these limitations and provide a more comprehensive view of the pandemic in Algeria, 449 complete sequences retrieved from the GISAID database were analyzed using different bioinformatics tools. This integrative approach helped us better understand how the virus evolved in Algeria, guiding future public health strategies.

## Materials and methods

### Sample collection and processing

Samples were obtained as part of the National SARS-CoV-2 Genomic and Variant Surveillance Initiative at the Pasteur Institute of Algeria (designated as the reference laboratory). Additionally, a real-time RT-PCR assay targeting the SARS-CoV-2 genes was conducted. For positive cases, sequencing was performed using an amplicon-based strategy, which produced 36 overlapping amplicons of 1,000 bp each through multiplex PCR. Sequencing was performed using the MinION platform. All sequences were deposited in the GISAID database and are freely available ^14^.

### Temporal signal assessment, time-calibrated phylogeny (temporal scaling phylogeny) and continuous phylogeographic analysis

In total, 334 Algerian sequences were retrieved from the GISAID database in December 2022. Sequence alignment was performed using the MAFFT web server with fast and rough FFT-NS-2 parameter ^15^. Then, a maximum likelihood tree was generated using IQ-TREE software, with the best substitution model selected as GTR+F+R2 ^16^. The tree file from IQ-TREE was used as input for TempEst to test the temporal signals. Regression analysis of root-to-tip genetic distances in relation to sampling times showed a strong positive correlation coefficient (r = 0.96 and R² = 0.92), indicating the suitability of the dataset for phylogenetic molecular clock analysis ^17^. Subsequently, BEAST v1.10.4, was used to generate a tip-dated phylogenetic tree. For the nucleotide substitution model, GTR + F was applied with a gamma distribution. For continuous phylogeography, the latitude and longitude of Algerian cities were combined with the Brownian random walk model. All analyses were conducted under a lognormal uncorrelated relaxed clock and the Bayesian skyline plot (12 groups were used) as the nonparametric coalescent model. The MCMC chains were run for 500 million generations, with sampling every 10,000 generations. Modifications were made to the priors, with the ucld mean and tree model root height set to follow an exponential distribution ^18^.

TRACER v1.6.0 was used to examine the effective sample size (ESS), which exceeded 150 ^19^. TreeAnnotator v1.10.4 was then applied to annotate the maximum clade credibility (MCC) trees, which were visualized using FigTree v1.4.4 ^20,21^. Finally, SpreaD3 v0.9.7 software was used for phylogeographic visualization ^22^.

### Genomic investigation

#### Mutation analysis

As of January 2024, additional sequences have been added to the GISAID database. A dataset comprising 449 complete Algerian genomes was analyzed using Nextclade to identify mutations specific to the viral strains circulating in Algeria and the clustering pattern of the Algerian sequences. Furthermore, a sub-dataset containing 192 Algerian sequences, along with the Wuhan reference sequence, was created by applying two filters: selecting complete genomes with more than 29,000 nucleotides and manually selecting sequences with less than 2% Ns. To evaluate the selective pressure at the protein level, SNAP v2.1.1 software was used. The ω ratio, representing the ratio of non-synonymous mutations to synonymous mutations, was calculated for each sequence pair across all SARS-CoV-2 genes: *ORF1a, ORF1b, S, ORF3a, E, M, ORF6, ORF7a, ORF7b, ORF8, N,* and *ORF10* ^23,24^.

#### Effects of mutations on SARS-CoV-2 proteins

The 449 Algerian sequences were individually analyzed using the CoV-server app in the GISAID database to identify mutational patterns and amino acid variations. PredictSNP was subsequently used to assess the functional impact of these changes on the corresponding proteins ^25^.

#### Recombination detection and analysis

Recombination events were evaluated using the recombination detection program (RDP5). This program analyzes nucleotide sequences by comparing their similarities to detect mosaic sequences that show statistically significant differences ^26^.

### Haplotype network analysis

To generate the haplotype file in Nexus format, DnaSP v6.12.03 software was used. POPART was then employed to create a graphical representation of the evolutionary relationships among haplotypes, resulting in a haplotype network diagram. The analysis was conducted using 192 complete genomes (no missing data) under the default settings (epsilon = 0) ^27^.

## Results

### Phylogenetic analysis

The estimated SARS-CoV-2 evolutionary rate of Algerian sequences was 1.43 × 10⁻³ [7.0516 × 10⁻⁴–1.454 × 10⁻³]. In contrast, phylogenetic analysis revealed substantial diversity among sequences, with distinct clustering patterns observed despite their geographic origin, highlighting multiple introductions. However, sequences from the same city were occasionally grouped, emphasizing local transmission. The inclusion of time as a variable in the analysis allowed for the visualization of the evolutionary dynamics over time (Figure 1).

**Figure 1:**
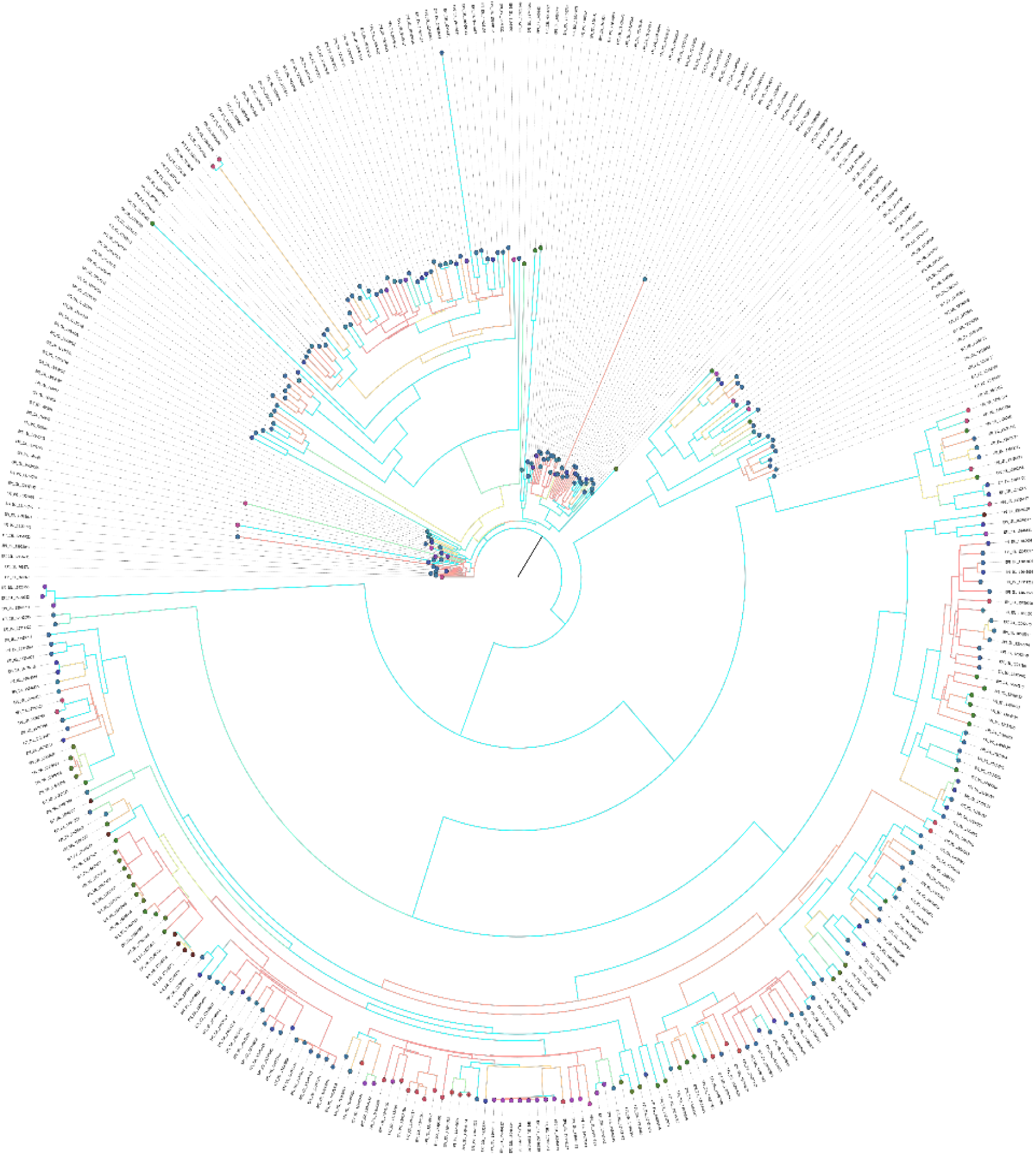
Bayesian phylogenetic tree based on a time-reversible model with unequal rates and unequal empirical base frequencies nucleotide substitution model (GTR+F), using BEAST. Colors indicate posterior, and the values are indicated at the end of each node: blue, high posterior probabilities; red, low posterior probabilities. Yellow indicates intermediate values.

### Diversity among Algerian sequences

Similarly, based on the Nextclade results displayed in Supplementary Figure 1, both multiple introductions and local transmission were observed. Sequences fell into different clades despite sharing a similar origin; however, in some cases, local clusters were identified in the phylogenetic trees. Furthermore, two sequences (highlighted in grey) were classified as part of a recombinant lineage.

***Supplementary figure 1:*** Classification of Algerian sequences in the Nextclade phylogenetic tree. The phylogeny was generated using Nextclade with GISAID as the database. A total of 449 Algerian sequences were introduced, and the colored dots indicate the clades to which each sequence belongs. The colors, specified in the legend, represent different clades, with grey indicating recombinant sequence.

### Phylogeographic diffusion in continuous space

To explore the spread of SARS-CoV-2, a continuous phylogeographic analysis was conducted. This approach is particularly useful for reconstructing transmission routes and identifying origins by estimating ancestral viral locations based on the latitudes and longitudes of the sampling sites. The differently colored nodes represent the sampled cities, and the blue nodes indicate internal nodes corresponding to ancestral locations (Figure 2).

**Figure 2:**
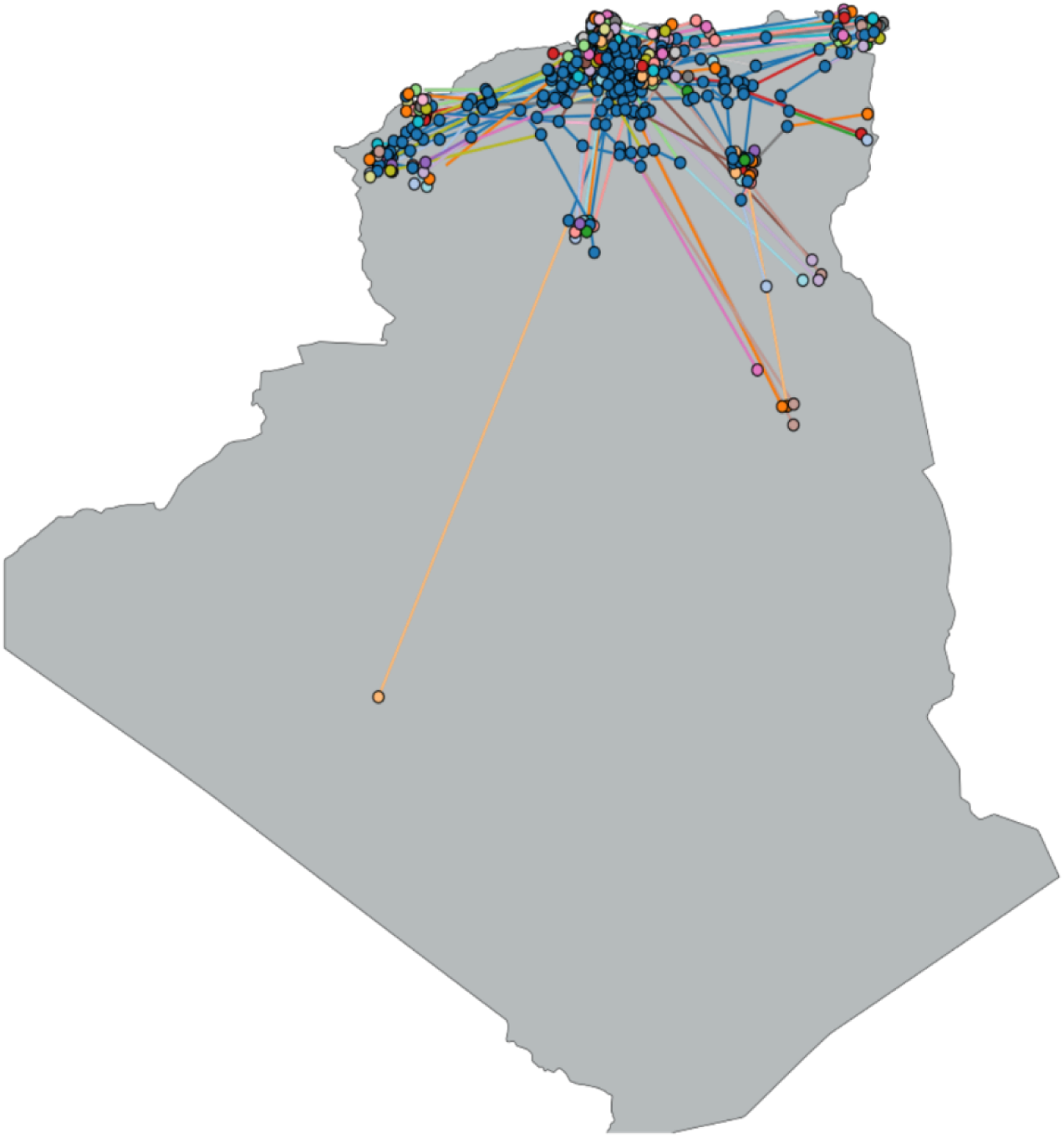
SARS-CoV-2 diffusion in continuous space in Algeria. The colored lines indicate the diffusion routes, the differently colored dots indicate the sampling locations, and the blue dots indicate the internal nodes.

### Phylodynamic analysis

The Bayesian skyline plot in Figure 3 provides insights into the population dynamics of SARS-CoV-2 over time, as inferred from the phylogenetic tree. The curve illustrates the changes in effective population size (Ne) over time. Peaks represent periods of population growth, valleys indicate periods of population decline, and the width of the curve reflects the uncertainty of the Ne estimates. As shown, there are three distinct peaks: the first in 2020, the second in 2021, and the last in 2022, corresponding to the different waves of the pandemic.

**Figure 3:**
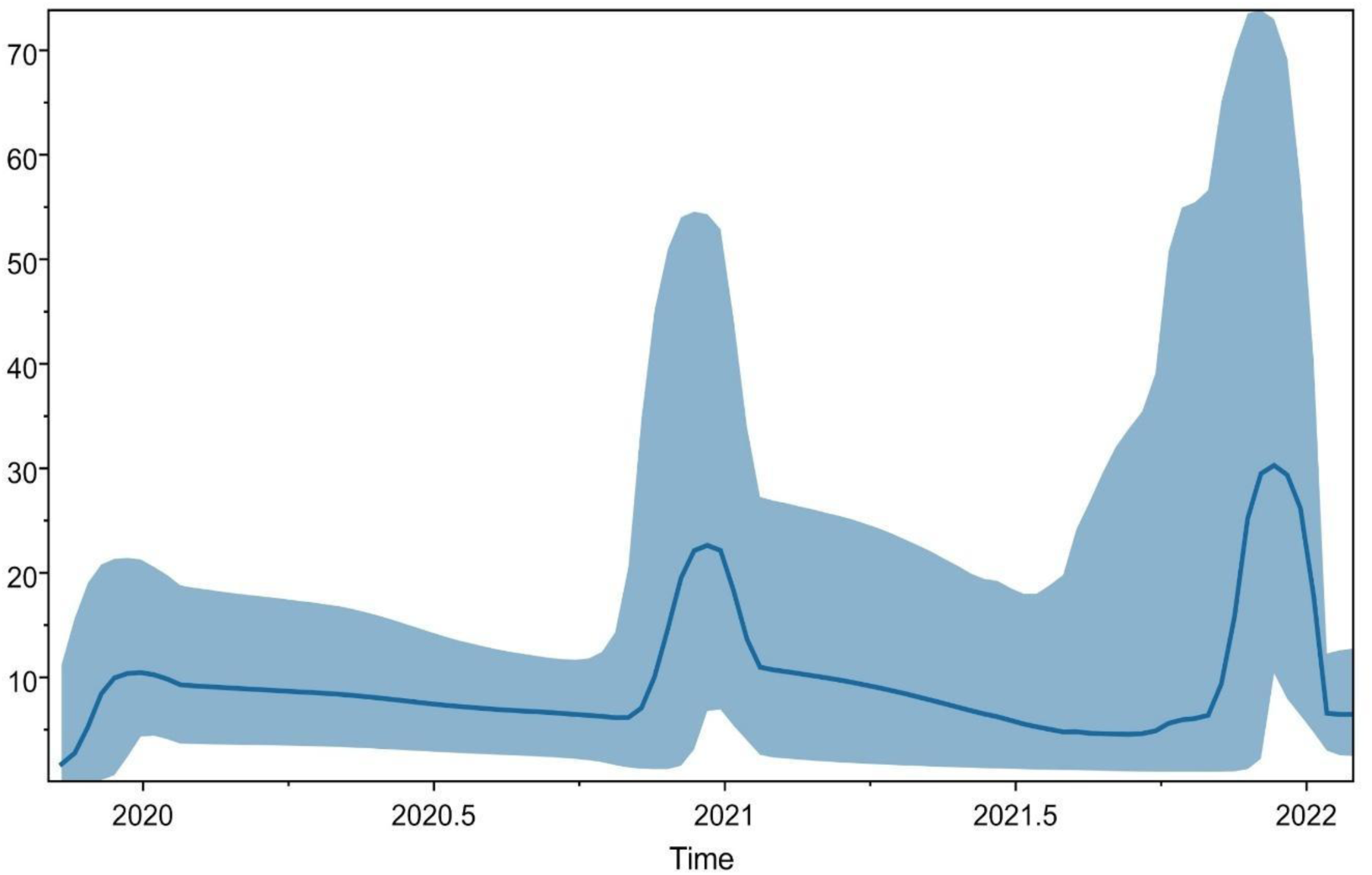
Bayesian skyline plot. Bayesian skyline plot of SARS-CoV-2 Algerian sequences illustrating changes in the effective population size over time.

### Genomic investigation

#### Selection pressure

The evolutionary selection pressure for each gene was evaluated using codon-by-codon cumulative behavior plots illustrating synonymous and nonsynonymous substitutions (**Supplementary figure 2**) and the calculated values of the nonsynonymous (dN) to synonymous (dS) mutation ratio (ω), summarized in Table 1. The ω ratio (dN/dS) indicates positive selection if it is greater than 1, and negative selection if it is less than 1.

**Table 1:**
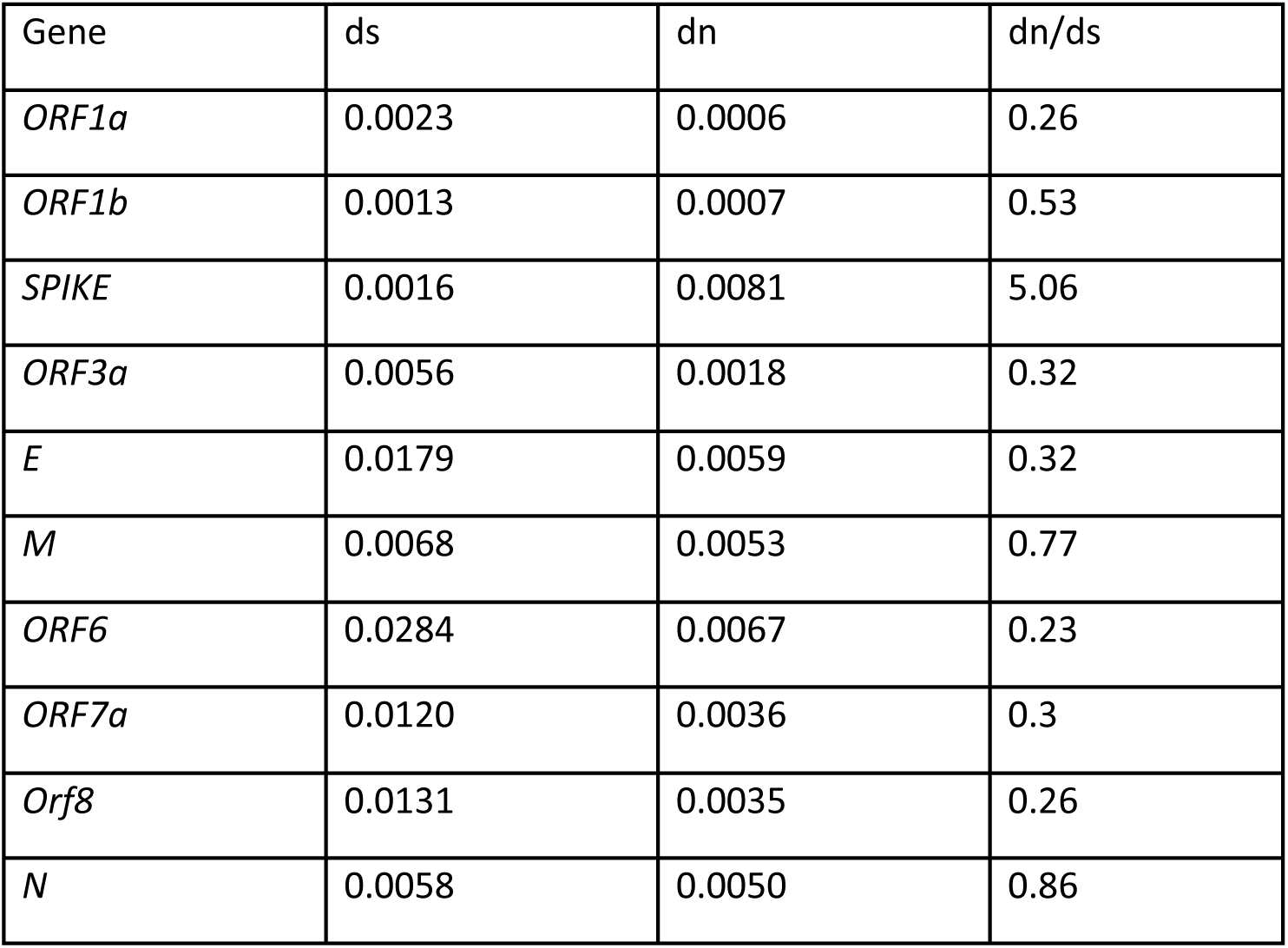
Selection Pressure of SARS-CoV-2 genes. A summary of the data generated by SNAP, assessing the selective pressure of each gene. Ds: observed synonymous substitutions; dn: observed nonsynonymous substitutions. dn/ds: selection pressure.

As shown in Table 1, *ORF1a* exhibited a highly negative selection with a value of 0.26, along with *ORF3a,* the *E gene, ORF6, ORF7a,* and *ORF8*. In contrast, *ORF1b*, the *M gene*, and the *N gene* displayed moderate negative selection. However, the *Spike gene* showed a value of 5.06, indicating strong positive selection. For *ORF10*, no significant results were observed.

The results from SNAP also included plots representing the insertion and deletion ratio (in/del), synonymous mutation rate (syn), and non-synonymous mutation rate (non-syn). Plots were generated for each gene.

***Supplementary figure 2:*** Plots of insertion/deletion, synonymous, and nonsynonymous mutation rates for each nucleotide in the genes. The plot illustrates how genetic changes (substitutions) occur in two specific types, synonymous and nonsynonymous, across each analyzed gene. The accumulation of amino acid substitutions in 198 Algerian sequences was examined, excluding the SARS-CoV-2 reference sequence from Wuhan, China. In the graph, the X-axis represents the codons, and the Y-axis represents the number of substitutions. The red line corresponds to synonymous substitutions, the green line represents nonsynonymous substitutions, and the gray vertical line marks stop codons.

As shown in supplementary figure 2, only *ORF1a* exhibited a higher in/del ratio than the synonymous and nonsynonymous mutation rates, whereas some other genes, such as *ORF1b*, *E, M, Spike*, and *ORF8*, showed a higher rate of non-synonymous mutations. In contrast, ORF3a, ORF7a, and ORF6 had a higher frequency of synonymous mutations than non-synonymous mutations.

In *ORF1a*, the in/del number was significantly higher than both the synonymous and non-synonymous mutation rates. In *ORF1b*, however, non-synonymous mutations predominated (**Supplementary figure 2**). Interestingly, the *Spike, E, M, N,* and *ORF8* genes all showed a higher rate of non-synonymous mutations than the other genes. In *ORF3a*, *ORF6*, and *ORF7a*, the synonymous mutation rate was higher than that of both in/dels and non-synonymous mutations. For *NSP10*, no significant results were observed.

#### Mutation analysis

Using PredictSNP, mutations present in Algerian sequences from the beginning of the Pandemic and until May 2023 were analyzed. The webserver tool integrates data from various mutation analysis methods (MAPP, PhD-SNP, PolyPhen-2, PolyPhen-1, SIFT, SNAP) (**Supplementary Table 1**) to provide a percentage indicating whether a mutation is deleterious (red) or neutral (green).

For NSP1, the mutation S135R, which has been present since the start of the pandemic, was classified as deleterious. In NSP2, the most common mutation was Q376K, which was neutral and exclusively found in 2022 sequences. In contrast, NSP3 and NSP6 mutations were recent, with each having one deleterious mutation, G489S and R233C, respectively, while the others were neutral. Similarly, mutations in NSP4, NSP5, NSP13, NSP14, NSP15, E, and NSP7b were neutral and occurred in many recent sequences. Spike mutations were neutral in both early and recent sequences. M and NS8 had one deleterious mutation each, I82T and R52I, respectively, along with additional neutral mutations (Supplementary Table 1). Within NSP12, the mutations T267M and N865H were identified as deleterious. Likewise, mutations in the NS3 protein, including T151I, Q57H, L129F, T223I, and G100C, negatively impacted the protein. Notably, mutations in the nucleocapsid N protein, such as S186Y, S194L, A251V, G204R, G204P, R203M, and S202N, exhibit deleterious effects on its structure and function.

**Supplementary Table 1:** Mutations in SARS-CoV-2 proteins (full set of Algerian sequences). Red indicates deleterious mutations, while green indicates neutral mutations.

Using CovSurver, several mutations specific to the Algerian sequences were identified. In NSP2, leucine (L) was substituted with valine (V) at position 270 in the sequence EPI_ISL_18090022. In NSP3, several mutations were identified, including a substitution of glycine (G) to alanine (A) at position 145 in the sequence EPI_ISL_15946147, phenylalanine (F) to tyrosine (Y) at position 210 in the sequence EPI_ISL_15928079, phenylalanine (F) to isoleucine (I) at position 336 in the sequence EPI_ISL_15235461, and alanine (A) to glycine (G) at position 1431 in the sequence EPI_ISL_12156740. In NSP6, leucine (L) was substituted with valine (V) at position 185 in the sequence EPI_ISL_15928087. In NSP14, a substitution of glycine (G) with serine (S) at position 189 was observed in sequence EPI_ISL_3375626. In NSP15, two mutations were detected: a substitution of lysine (K) to isoleucine (I) at position 307 in the sequence EPI_ISL_15928152 and tryptophan (W) to lysine (K) at position 332 in the sequence EPI_ISL_12156748 (Table 2).

**Table 2:** Mutations exclusive to Algerian SARS-COV-2 sequences. A summary of the Predict SNP results of the classification of Algerian SARS-CoV-2 specific mutations.

| Protein | Mutations | PredictS NP | MAPP | Phd-SNP | PolyPhe n-1 | PolyPhe n-2 | SIFT | SNAP |
| --- | --- | --- | --- | --- | --- | --- | --- | --- |
| NSP2 | L270V | 63% | 84% | 83% | 67% | 63% | 76% | 58% |
| NSP3 | G145A | 63% | 65% | 72% | 67% | 68% | 79% | 50% |
| NSP3 | F210Y | 83% | 77% | 68% | 67% | 63% | 71% | 50% |
| NSP3 | F336I | 76% | 65% | 73% | 59% | 81% | 79% | 81% |
| NSP3 | A1431G | 83% | 73% | 83% | 67% | 61% | 61% | 50% |
| NSP6 | L185V | 75% | 64% | 78% | 67% | 63% | 76% | 71% |
| NSP14 | G189S | 74% | 75% | 72% | 67% | 41% | 76% | 67% |
| NSP15 | K307I | 87% | 86% | 73% | 74% | 54% | 79% | 81% |
| NSP15 | W332K | 87% | 92% | 88% | 74% | 81% | 79% | 89% |

#### Recombination detection and analysis

Recombination analysis revealed one recombination event with strong statistical support. The recombinant sequence was detected by seven methods out of nine, with a significant p-value. The results are summarized in Table 3. The sequence in question is EPI_ISL_15920753 sampled from a Male on the 25^th^ of September 2022 in the city of Sétif and belonging to the BA.2 sub-lineage of the omicron variant. The Major parent, which by definition is the genetically closer parent to the recombinant sequence and contributes to a larger portion of the genome, is EPI_ISL_15790700 sampled on the 18^th^ of August 2022 in the city of El_Taref and belonging to the omicron variant sub-lineage BA.2.38. The minor parent was EPI_ISL_12156732 sampled on the 1^st^ of October 2020 from Blida and belonging to the B.1.1 lineage from the early pandemic.

**Table 3:**
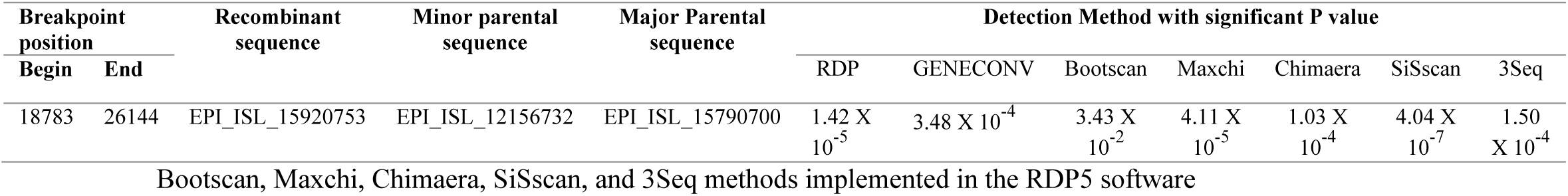
Recombination analysis of 193 high-quality full Algerian genomes using RDP, GENECONV.

### Haplotype network analysis

Haplotype analysis revealed 179 distinct haplotypes, highlighting the extensive diversity within our dataset (**Figure 4**). Colored nodes represent the Algerian cities where the samples originated, and black nodes denote internodes, indicating missing or unsampled data. Sequences from the same location can cluster together, although some diverge significantly despite sharing the same origin. Interestingly, a potential superspreader event can be identified based on the central and highly connected nodes observed around H91 in the abovemented figure. The identification of haplotype H91 in distant cities, combined with several directly derived haplotypes collected within a brief time frame, strongly indicates a superspreader event. This could be a result of a highly mobile infected person combined with a significant gathering that promoted extensive spread of the virus.

**Figure 4:**
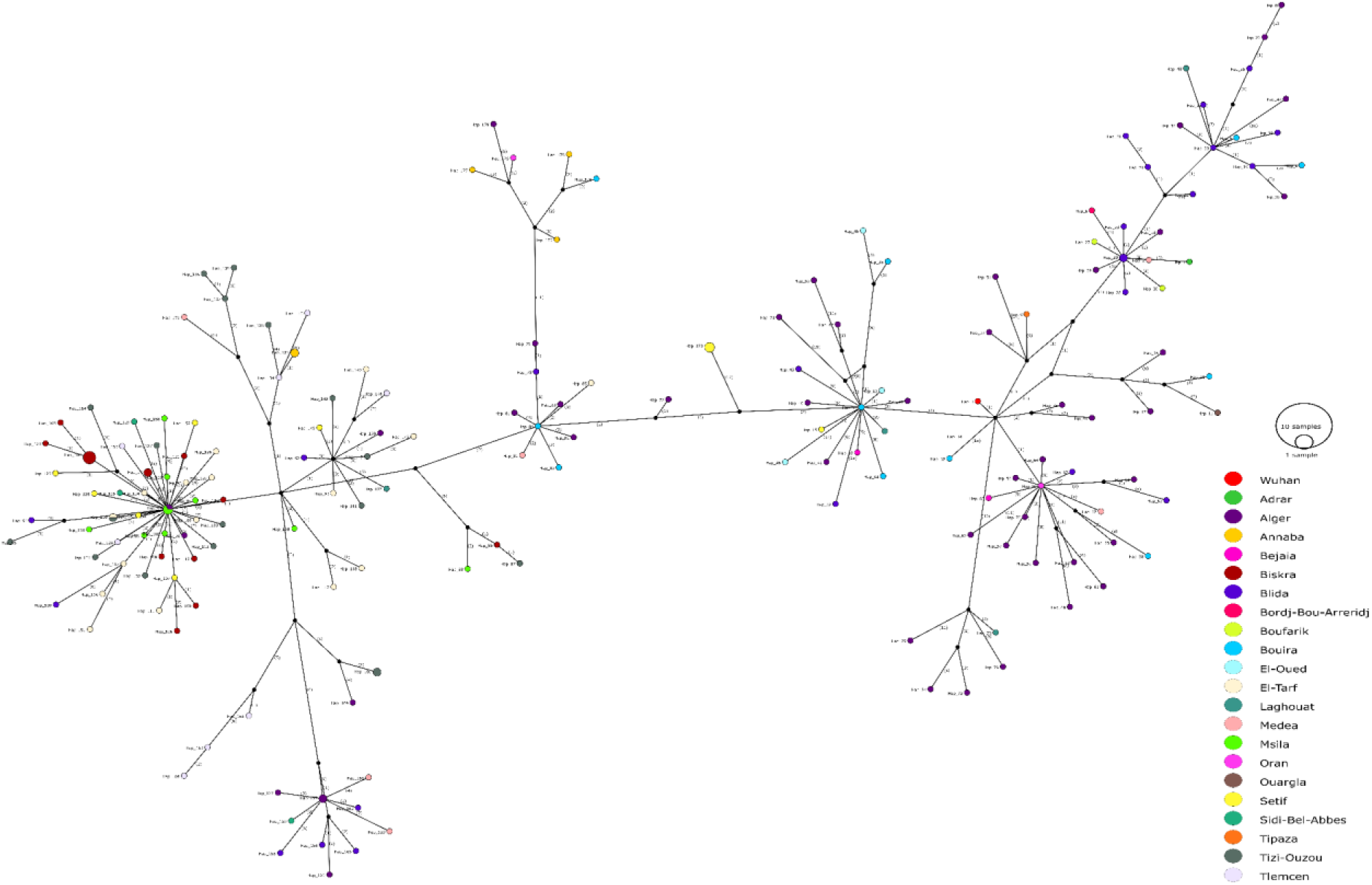
Haplotype network plot of SARS-CoV-2 Algerian sequences. Different colors indicate the sampling locations. The black circles indicate internal nodes pointing to missing unsampled data. The size of the circles is proportional to the number of sequences in each cluster.

## Discussion

Exploring the evolution of SARS-CoV-2 remains critical for understanding viral adaptation over time and for leading public health interventions. As the virus continues to circulate globally, selective pressures such as human immunity and public health interventions trigger its evolutionary changes and adaptations. Hence, genomic surveillance enables experts to anticipate shifts in transmissibility or virulence ^28^. It plays a key role in the development of vaccines. For instance, mutations in the spike (S) protein, especially in the receptor-binding domain, might reduce the efficacy of already existing vaccines, henceforth requiring periodic updates to hold protection against new emerging variants ^29^. Likewise, it increases the reliability of diagnostic tools by identifying mutations that may alter test sensitivity ^30^. In parallel, the real-time identification of new variants is crucial for guiding mitigation strategies, such as travel restrictions and targeted lockdowns, for effective outbreak prevention and control ^31^. To explore this virus more efficiently, instead of directly measuring the effects through laboratory experiments, which is challenging due to the lack of practical assays for many viral proteins, the approach is to use vast publicly available SARS-CoV-2 genetic sequences to estimate the effects of mutations ^32^. This is the same approach that we used, focusing only on Algerian sequences.

Our study presents a comprehensive analysis of 449 SARS-CoV-2 sequences from Algeria, including a phylogenetic tree, continuous-space phylogeographic diffusion, phylodynamic analysis, recombination analysis, selection pressure on genes, specific mutation analysis, and haplotype network analysis. First, when comparing the current mutation rate (1.43 × 10⁻³) to previous estimations, including 18 full genomes and 11 partial sequences derived from Algeria (5.4043 × 10−4) and performed at the beginning of the pandemic, a significant increase is clearly observed ^13^. This is in line with what is expected for fitness improvement and enhancement ^33^.

Furthermore, the phylogenetic tree results showed diverse and distinct clades in the dataset, indicating multiple introductions of the virus into Algeria. This was already observed at the beginning of the pandemic when travel restrictions were very strict, and the dataset was smaller ^13^. Strikingly, with a larger dataset, different sequence clustering and tree branching were observed in this study. For example, the sequence EPI_ISL_766862 from Bouira was previously clustered with EPI_ISL_766866 from the same location (posterior probability = 80%) ^13^. In the new analysis, it branched with EPI_ISL_15928087 from Algiers (posterior probability = 100%), which was sampled on the 13^th^ of October 2020 and submitted to the GISAID database on the 28^th^ of November 2022. This highlights the importance of sampling efforts, sequencing, and real-time reporting in genomic epidemiology. Similarly, sequences from different cities clustered together, highlighting the impact of local travel on the spread of the virus. However, local transmissions at the city level were clearly observed, indicating the impact of gatherings on the spread of the virus. This occurred alongside the ease or discontinuation of mitigation measures. This is in accordance with the behavioral analysis during the COVID pandemic in Algeria, Tunisia, and Morocco ^34^.

Similarly, the phylogeographic diffusion analysis in continuous space allowed us to visualize the spread of the virus over time and trace back the ancestral locations. Notably, most ancestral locations were in the northern and central parts of Algeria; in the South, only one ancestral location was observed. This pattern correlates with the high population density and travel connectivity in these regions, as well as the sampling bias, where northern and central Algeria have better genomic surveillance than the southern part of the country ^35,36^. As expected, the Bayesian skyline plot demonstrated three distinct peaks for population size growth, which reflects the different waves resulting from the appearance of new variants ^37^.

When evaluating the selection pressure among the different structural, nonstructural, and accessory genes, the *S gene* had a significant increase in the ω value compared to the previous Algerian study ^13^. Therefore, the impact of larger datasets in providing more statistical power to detect positive or purifying selection is undoubtedly highlighted in this study. Nonetheless, most sequences were sampled from 2022 and 2023; consequently, the results were influenced by spatiotemporal bias. This manifests as higher ω ratios in the *M* (0.77) and *N* (0.86) genes, exceeding typical global estimates (0.1-0.2 for *M* and 0.2-0.4 for *N*) ^38,39^. Hence, reflecting variant-specific adaptation. During 2022–2023, delta and omicron lineages predominated with mutations like I82T in M and R203K/G204R in N proteins, formerly associated with changes in viral replication, assembly, and immune evasion ^40^. Moreover, since the dataset is mainly composed of recent sequences from a limited geographic region, this potentially influences the reported results. This bias may lead to increased ω values by capturing current founder effects, selective sweeps, or transitory polymorphisms rather than long-term purifying selection pressure ^41^. Therefore, our results emphasize the need to consider both sampling time and geographical origin when interpreting ω values, as local epidemiological factors and host immune pressures might prompt distinct adaptive trajectories in different viral populations.

For the mutation analysis, a few deleterious mutations were recognized. The mutation T151I in NS3 (ORF3a) was found to lead to structural damage, specifically a buried H-bond breakage in both Alpha and Beta variants, resulting in a loss-of-function mutation ^42^. Additionally, the mutation Q57H was found to cause significant structural changes in the dimer conformation ^43^. This could lead us to conclude that the other two mutations, L129F and G100C, may have a similar effect. Moreover, S194L and S202N identified in N (ORF9) were found to be two effective mutations ^44^. Further, the deleterious effects of mutations T267M and N865H, as well as S186Y, A251V, G204R, and S202N in NSP12 and N, respectively, were revealed. One important mutation found was R233C in NSP6, which was demonstrated to be deleterious. It was also identified as a missense mutation, altering the NSP6 protein function ^45^. Additionally, the mutation T223I, found only in recent Algerian sequences and having a deleterious effect, was associated with less efficient replication and potentially contributes to lower pathogenesis in the Omicron strain ^46^. More notably, the analysis of protein sequence mutations revealed nine mutations unique to the Algerian SARS-CoV-2 dataset. Among these, three mutations were detected to have deleterious effects using Predict-SNP: F336I in NSP3, and K307I and W332K in NSP15. Each of these mutations was identified in a single sequence, EPI_ISL_15235461, EPI_ISL_15928152, and EPI_ISL_12156748, respectively. It was revealed that NSP15 plays a crucial role in host RNA processing, and mutations within this protein may alter viral replication dynamics, potentially affecting viral fitness ^47^. Also, studies show that a mutation in NSP3 (S676T) can reduce viral replication by disrupting its interaction with the host’s translation factors, however, a compensatory mutation in the Nucleocapsid protein (N-S194L) can restore the virulent activity ^48^. This emphasizes the functional role of NSP3 and its interaction with other proteins in modulating SARS-CoV-2 replication. Remarkably, these mutations have not been previously reported in the scientific literature, underscoring the need for further investigation into their functional implications. While some of these mutations may later appear in sequences worldwide, this initial identification shows the importance of regional genomic surveillance in capturing emerging viral variations.

Strikingly, based on the Nextclade analysis, two recombinant sequences were present in the dataset. The EPI_ISL_12043095 sampled, in February 2022 from a 76-year-old male, was first classified by the Pangolin software as BA.2 omicron subclade. However, when conducting this new investigation, it turned out to be part of the XAP recombinant clade. This is a key observation as it predates the official designation of XAP by two months, suggesting earlier cryptic circulation ^49^. Similarly, the sequence EPI_ISL_17985292 was collected in April 2023 and belongs to the XBV recombinant clade, which was first reported in March 2023 according to the Outbreak.info reporting system. Nonetheless, according to the GISAID database, it was already circulating in the USA on the 14^th^ of November, 2022 ^50,51^. Furthermore, the RDP 5 software detected a third recombinant sequence EPI_ISL_15920753 from 2022 involving a major parent form sublineage BA.2.38, and a minor parent from B.1.1 sublineage. This might be explained by several hypotheses, such as undetected persistence where the B.1.1 lineage or its close relative were still present in under-sampled populations or immunocompromised individuals, hence making them less likely to be detected in genomic surveillance at the time of the recombination (2022). Similarly, the ancestor proxy effect, where the B.1.1 is not the actual minor parent. Instead, it is the closest known lineage to the actual unsampled parental sequence ^52,53^.

All the results above demonstrate the diversity and high mutation rate of SARS-CoV-2. From previous studies conducted at the beginning of the pandemic, there was not much variation, especially in non-structural genes. However, for the Spike gene, the mutation rate was high as early as April 2020 and increased even further by January 2023.

We cannot deny that the number of samples is very important for viral evolutionary studies. With our dataset comprising 449 sequences, we were able to obtain valuable insights. However, additional data is still needed for more precise results. Previous studies from Algeria had approximately only 18 fully sequenced genomes, while ours contained 193 full, high-quality sequences (no gaps or n stretches). With this amount of data, far more information on Algerian lineages were obtained, and 197 haplotypes were described within a single country ^13^. Two types of clusters Were obtained when performing the haplotype network analysis clusters from the same provenance and others from completely different sampling locations. This demonstrates how lifted restrictions impacted the virus’s spread, increased its mutation rate, and amplified its diversity.

Note that the evolution of SARS-CoV-2 is very dynamic; for instance, mutations that were identified as Algerian characteristic signatures are no longer classified as such due to the growing number of samples and data. Back in 2020, the amino acid replacement A130V in the NSP12 gene was first classified as an Algerian characteristic mutation; however, in recent analyses, it was demonstrated to originate in Abu Dhabi (EPI_ISL_69815) on 2020-06-12. Thereafter, it was imported to Algeria (EPI_ISL_766874) on 21/06/2020. On the other hand, new Algerian substitutions arose, namely the W332K mutation, which was first reported in Algeria on 14/03/2021. This demonstrates the importance of molecular investigation in tracking new mutations that might impact drug and vaccine development, as well as mitigation measures. Moreover, it enables molecular tracing. For instance, the aforementioned mutation was later spotted in the UK on 26/06/2022. Similarly, the N874H mutation in NSP12, found in EPI-ISL-766875, was determined to have a deleterious effect. In the latest analysis, it was found to have migrated first to the UK on 06/07/2020; however, we demonstrate that its first appearance was actually in Libya on 18/06/2020, then in Egypt on 01/07/2020, and later in the UK. Despite the physical barriers and lockdown measures, the spillover of the virus within bordering countries can clearly be witnessed.

Finally, the haplotype network analysis of Algerian SARS-CoV-2 sequences demonstrated a heterogeneous pattern of viral transmission, with several individual clusters and highly connected central nodes. Particularly, a remarkable cluster centered on a haplotype associated with Algiers and M’sila. It displayed a star-like structure, in which various sequences radiate from a single central node with a small number of mutations. This topology is distinctive of a superspreader event, where a single infected individual or a small number of individuals engender a large number of secondary cases within a brief timeframe ^54,55^. Such events can increase transmission, reduce short-term genetic diversity, and speed up the spread of specific viral variants ^56^. These dynamics are evident in the recorded low genetic divergence and restricted mutational branching within the cluster, concordant with a rapid dissemination from a single founder. On the other hand, the existence of shared haplotypes across multiple cities suggests inter-regional movement and transmission. This emphasizes the role of human mobility and superspreading in shaping viral population structure.

## Conclusion

Overall, our in-depth analysis focused on the evolutionary dynamics of the Algerian SARS-CoV-2 pandemic. Hence, helping to fill a critical knowledge gap. It emphasizes the importance of real time genomic surveillance and reporting in supporting decision-making, evaluating diagnostic tools, and guiding therapeutic and vaccination strategies. Additionally, it underlined the massive impact of the dataset size on the results and conclusions to be drawn.

## Acknowledgments

We acknowledge the Pasteur Institute of Algeria for providing the sequences in the GISAID’s EpiFlu database, which formed the foundation for this research.

